# A Machine Learning Approach to Map Tropical Selective Logging

**DOI:** 10.1101/451856

**Authors:** MG Hethcoat, DP Edwards, JMB Carreiras, RG Bryant, FM França, S Quegan

**Affiliations:** School of Mathematics and Statistics, University of Sheffield, Sheffield, S3 7RH, UK; Grantham Centre for Sustainable Futures, University of Sheffield, Sheffield S10 2TN, UK; Department of Animal and Plant Sciences, University of Sheffield, Sheffield S10 2TN, UK; National Centre for Earth Observation, University of Sheffield, Sheffield, S3 7RH, UK; Department of Geography, University of Sheffield, Sheffield, S3 7ND, UK; Lancaster Environment Centre, University of Lancaster, Lancaster, LA1 4YQ, UK; Instituto de Ciências Biológicas, Universidade Federal do Pará, Belém, 66075-110, Brazil.

**Keywords:** Brazil, Conservation, Degradation, Landsat, Random Forest, Selective logging, Surface reflectance, Texture measures, Tropical forests

## Abstract

Hundreds of millions of hectares of tropical forest have been selectively logged, either legally or illegally. Methods for detecting and monitoring tropical selective logging using satellite data are at an early stage, with current methods only able to detect more intensive timber harvest (>20 m^3^ ha^−1^). The spatial resolution of widely available datasets, like Landsat, have previously been considered too coarse to measure the subtle changes in forests associated with less intensive selective logging, yet most present-day logging is at low intensity. We utilized a detailed selective logging dataset from over 11,000 ha of forest in Rondônia, southern Brazilian Amazon, to develop a Random Forest machine-learning algorithm for detecting low-intensity selective logging (< 15 m^3^ ha^−1^). We show that Landsat imagery acquired before the cessation of logging activities (i.e. the final cloud-free image of the dry season during logging) was better at detecting selective logging than imagery acquired at the start of the following dry season (i.e. the first cloud-free image of the next dry season). Within our study area the detection rate of logged pixels was approximately 90% (with roughly 20% commission and 8% omission error rates) and approximately 40% of the area inside low-intensity selective logging tracts were labelled as logged. Application of the algorithm to 6152 ha of selectively logged forest at a second site in Pará, northeast Brazilian Amazon, resulted in the detection of 2316 ha (38%) of selective logging (with 20% commission and 7% omission error rates). This suggests that our method can detect low-intensity selective logging across large areas of the Amazon. It is thus an important step forward in developing systems for detecting selective logging pan-tropically with freely available data sets, and has key implications for monitoring logging and implementing carbon-based payments for ecosystem service schemes.

## 1. Introduction

Earth’s tropical forests are being rapidly lost and degraded by agricultural expansion and commercial logging operations, with population growth projected to further increase pressures on forests globally (Asner et al., 2005; DeFries et al., 2010). The ability to monitor forest disturbances is an important component in sustainable forest management, understanding the global carbon budget, and implementing climate policy initiatives, such as the United Nation’s (UN) Reducing Emissions from Deforestation and Forest Degradation (REDD+) programme, which seeks to mitigate climate change and biodiversity losses through improved forest management practices (GOFC-GOLD, 2016). The UN anticipates that payments to nations under REDD+ initiatives, which compensate countries for conserving forests (and sequestering carbon), could reach $30 billion annually (Phelps et al., 2010, UN-REDD Programme, http://www.un-redd.org)

Remote sensing is considered the most accurate and cost-effective way to systematically monitor forests at large scales (Achard et al., 2007; Herold and Johns, 2007; Shimabukuro et al., 2014). Large-scale monitoring of deforestation has significantly improved in recent years, and forest losses can be identified with accuracies greater than 90% using freely available satellite data (Hansen et al., 2013). In addition, near real-time deforestation tracking and alert systems are now possible with systems like DETER (Shimabukuro et al., 2012), FORMA (Hammer et al., 2014; Hansen et al., 2013), and Global Forest Watch (Hansen et al., 2016). In contrast, methods for detecting and monitoring forest degradation are less developed. Forest degradation is an ambiguous term, with over 50 different definitions and no internationally established description (Ghazoul et al., 2015; Simula, 2009). This makes generalizing its impacts difficult, in part because degradation can include forests subject to varying intensities of selective logging, fire, artisanal gold mining, fuelwood extraction, etc., which has hampered the development of coordinated international forest policies to track and monitor forest degradation (Ghazoul et al., 2015; Sasaki and Putz, 2009).

Here we focus on detecting a key driver of forest degradation globally, commercial logging operations. In contrast to forest clearance (i.e. deforestation), selective logging represents a more diffuse disturbance wherein only a subset of trees (typically the most economically valuable) are harvested (Fisher et al., 2014; Putz et al., 2001). The resulting forest maintains some degree of its original composition (e.g. canopy cover, biodiversity measures, carbon content, etc.) but is punctured by treefall gaps and logging roads and consequently lies on a continuum between primary forest and complete deforestation (Ghazoul et al., 2015; Thompson et al., 2013). The intensity of selective logging operations can vary in two main ways: (1) the volume of wood harvested typically ranges up about 50 m^3^ ha^−1^, as high as 150 m^3^ ha^−1^ in Asia (Burivalova et al., 2014; Putz et al., 2001) and (2) the degree to which reduced-impact logging is practiced, in which damage to the remaining forest is minimized by careful planning of road networks, skid trails, and directional felling of trees to limit additional tree or canopy damage (Putz and Pinard, 1993).

Selective logging activities are often the first anthropogenic disturbance to affect primary tropical forests (Asner et al., 2009b; Nepstad et al., 1999) and are thought to be a major source of carbon emissions from degradation (Hosonuma et al., 2012; Pearson et al., 2017). Moreover, road networks associated with logging are often precursors to additional land-use changes (such as agricultural conversion or development of human settlements) and facilitate further degradation (e.g. increased susceptibility to fires or illegal logging) and forest losses (Alamgir et al., 2017; Kumar et al., 2014; Matricardi et al., 2010). Estimates suggest over 400 million ha of tropical forest, an area the size of the European Union, are earmarked in the tropical timber estate to be logged (Blaser et al., 2011). However, the extent of forest subjected to selective logging across the tropics has yet to be estimated (Asner et al., 2005).

Several authors have tried to address the challenges of using satellite data to estimate forest disturbances from selective logging in the tropics (Asner et al., 2005, 2004a, 2002, Matricardi et al., 2010, 2007; Shimabukuro et al., 2014; Souza and Barreto, 2000; Souza et al., 2005). The majority of approaches employ classification of fractional images derived from spectral unmixing of Landsat scenes. Despite these advancements, Landsat imagery has been considered too coarse to monitor less intensive selective logging activities, with nearly all applications involving logging intensities > 20 m^3^ ha^−1^ (Asner et al., 2005, 2004a, 2002, Matricardi et al., 2010, 2007; Shimabukuro et al., 2014; Souza and Barreto, 2000; Souza et al., 2005). While most authors acknowledge their methods can detect areas of selective logging at moderately high intensities (> 20 m^3^ ha^−1^; 3-7 trees ha^−1^), that possess large canopy gaps and an abundance of spectrally distinct features, like log landing decks or large road networks, their respective abilities to detect lower logging intensities are unknown. Therefore, using Landsat data to map and quantify selective logging at lower logging intensities (< 20 m^3^ ha^−1^) remains a major challenge, and the amount of forest disturbance overlooked using currently available techniques is unknown. Yet, growing concerns over the impacts of selective logging on carbon and biodiversity (Bicknell et al., 2014; Edwards et al., 2014; França et al., 2017; Martin et al., 2015; Putz et al., 2008) has led to increased use of improved forest management practices, such as reduced-impact logging (Putz and Pinard, 1993). Consequently, the extent of tropical forests being logged at lower intensities and with reduced-impact is almost certainly expanding. In addition, there is an ever-increasing need to detect and account for the estimated 50-90% of tropical timber on the international market harvested illegally at very low intensities (Brancalion et al., 2018; Kleinschmit et al., 2016). Therefore methods to detect subtle forest disturbances from satellite systems with regular global coverage are urgently needed, both to establish reference levels from historical data (e.g. the vast amount of freely available Landsat archives) and to obtain maximum benefit from current and future systems, such as Landsat 8, 9 and Sentinel-2 (Drusch et al., 2012; Roy et al., 2014).

The primary objective of this study was to develop a new method for detecting selective logging in moist tropical forest with Landsat data. It focuses on reduced-impact selective logging of intensity < 15 m^3^ ha^−1^ (1-2 trees ha^−1^), much lower than is typically reported in studies that use remote sensing data to estimate selective logging (Asner et al., 2005, 2004a; Souza and Roberts, 2005), but still more than three times the background rate of natural mortality estimated for tropical forests (Brienen et al., 2015; Clark et al., 2004). We used detailed spatial and temporal logging records from Rondônia, Brazil, together with Landsat data, to build a machine learning algorithm for detecting selectively logged Landsat pixels. Machine learning (neural networks, decision trees, support vector machines, etc.) for classification of satellite imagery has been used with increasing success in recent years (Tuia et al., 2011) and can turn a suite of predictor variables weakly correlated with a response into a relatively strong classifier (Breiman, 2001). The successful application of this algorithm to a test site in northern Pará, Brazil, approximately 1500 km from the location of algorithm development, demonstrates that this approach is transferable and can greatly improve existing methods of detecting subtle selective logging activities in the tropics.

## 2. Study sites and satellite imagery

Data from two test sites in the Brazilian Amazon were used in this study (Fig. 1a). The Jamari site consists of *terra firme* tropical forest inside the Jamari National Forest, Rondônia, Brazil. The logging concession was subdivided into forest management units (FMUs) that were each approximately 2,000 ha (Fig. 1b). Selective logging occurred within a single FMU in each year, at an intensity of approximately 10 m^3^ ha^−1^ (1-2 trees ha^−1^), beginning at the end of the wet season (roughly June) and continuing through the dry season (until November) from 2011 through 2015. Forest inventory measurements were recorded by trained foresters and included the spatial location of each marketable tree species within the concession, its height, diameter, estimated volume, and if it was logged (only trees > 50 cm in diameter can be harvested). Overall, the spatial locations of more than 13,000 individually identified trees that were selectively logged between 2011 and 2015 were recorded. A field survey in 2016 relocated a subset of trees (*n* = 214) to estimate the geolocation precision of the logging inventory records (mean = 6.2 m; standard deviation = 6.6 m). This detailed record of where and when trees were selectively removed provided the means to build the machine learning algorithms described in Section 3.3 and to test their performance.

**Fig. 1.**
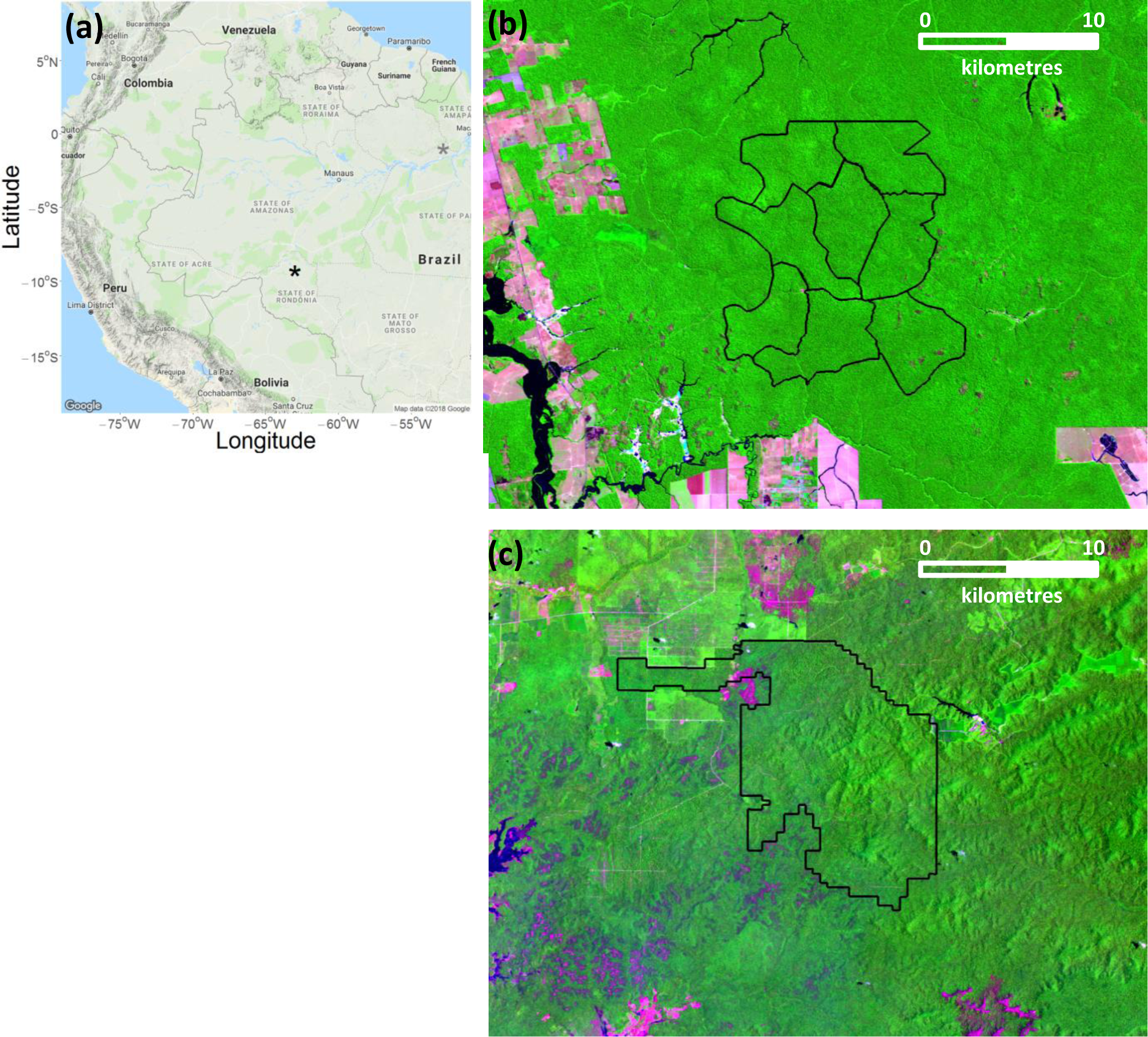
Location of the Jamari (black star) and Jari (grey star) study sites in the Brazilian Amazon (a). Landsat 8 image (RGB bands 6,5,4) of the Jamari site (b) from June 2016 in Rondônia, Brazil. The six southern forest management units (outlined in black) include the locations of data inputs for machine learning algorithm development, while the northern 2 units remained unlogged. Landsat 8 image (bands 6,5,4) of the Jari site (c) from September 2016 in Pará, Brazil. Jamari and Jari were selectively logged from 2011-2015 and in 2012, respectively.

At the Jamari site, heavy cloud cover typically occurs between October and May, but cloud-free images from Landsat 5 Thematic Mapper (TM), Landsat 7 Enhanced Thematic Mapper (ETM+), and Landsat 8 Operational Land Imager (OLI) were acquired approximately annually for 2008 to 2016 in the intervening dry season (Table 1). Note that the 2012 ETM+ images suffered from missing data as a result of the scanline corrector error and appear striped (Storey et al., 2005). For the analyses, we distinguished “early” and “late” images for a given region. The early image was the last cloud-free image of the dry season in the *same year* the FMU was logged (typically in August, approximately 2-3 months before cessation of logging activities for the season). The late image was the first cloud-free image of the dry season in the *year after* cessation of logging activities (typically in June, approximately 8-12 months after the FMU was logged). We used early and late imagery to generate two separate datasets and build two separate algorithms in order to assess which time period provided better detection of selective logging. This is illustrated for a hypothetical logging season in Fig. 2. The selection of two time periods reflects the fact that after 8-12 months, regrowth of foliage and other vegetation can reduce the spectral signatures required to identify canopy gaps and woody debris in tropical systems (Asner et al., 2004a, 2004b; Broadbent et al., 2006).

**Fig. 2.**
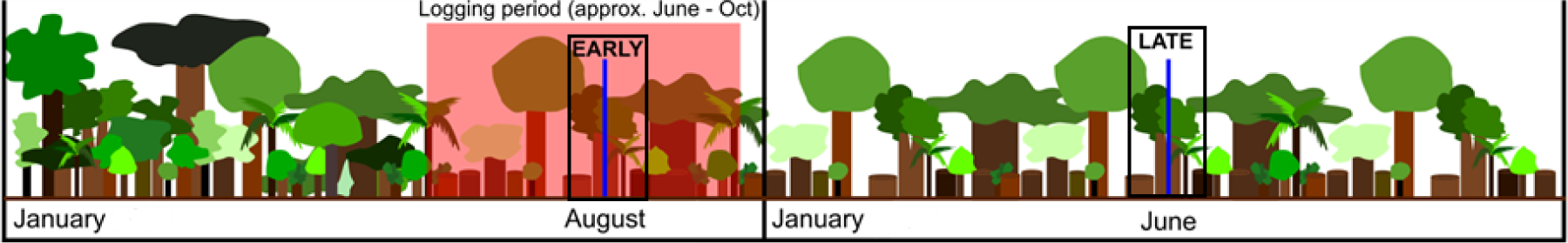
Timeline representation of a single forest management unit in the Jamari study site. Vertical blue lines indicate image acquisitions during the early and late time periods (black boxes) relative to when logging occurred (red box). In this example the early Landsat image was acquired part way through the logging season, so part of the management unit has yet to be cut. The late image is the first cloud-free image of the following dry season and is acquired approximately 8 months after the management unit was selectively logged.

**Table 1.** Landsat 5 (TM), 7 (ETM+), and 8 (OLI) scenes used to build and assess Random Forest models developed to detect selective logging. The Jamari study site is path 232, row 066 and the Jari site is path 226, row 061.

| Study Site | Acquisition Date | Scene Timing | Solar Zenith Angle | Landsat Sensor |
| --- | --- | --- | --- | --- |
| Jamari | 2008-07-28 | Early | 49.75 | TM |
|  | 2009-07-31 | Early | 50.00 | TM |
|  | 2010-07-18 | Early | 46.36 | TM |
|  | 2011-08-06 | Early | 51.67 | TM |
|  | 2012-08-16 | Early | 54.05 | ETM+ |
|  | 2013-08-27 | Early | 57.07 | OLI |
|  | 2014-08-30 | Early | 58.84 | OLI |
|  | 2015-09-02 | Early | 60.19 | OLI |
|  | 2009-06-29 | Late | 43.79 | TM |
|  | 2010-07-02 | Late | 43.63 | TM |
|  | 2011-07-05 | Late | 44.30 | TM |
|  | 2012-06-13 | Late | 41.64 | ETM+ |
|  | 2013-07-10 | Late | 42.26 | OLI |
|  | 2014-06-11 | Late | 40.43 | OLI |
|  | 2015-06-14 | Late | 40.47 | OLI |
|  | 2016-06-16 | Late | 40.37 | OLI |
| Jari | 2011-11-08 | Early | 123.31 | ETM+ |
|  | 2012-11-10 | Early | 125.27 | ETM+ |
|  | 2011-07-03 | Late | 48.12 | ETM+ |
|  | 2013-08-17 | Late | 60.92 | OLI |

The Jari site (Fig. 1c) in Pará, Brazil, consists of *terra firme* tropical forest inside the 12,500 ha Jari concession that was selectively logged at an intensity of approximately 12 m^3^ ha^−1^ (1-3 trees ha^−1^) between July and December 2012. In contrast to Jamari, the Jari site lacked detailed information on where trees were removed, but the volume of wood (m^3^) removed was recorded for 10 ha (400 m x 250 m) blocks in the concession. The Jari site allowed us to assess whether the algorithms developed using the Jamari dataset, located approximately 1500 km away, were transferable to this distant site. At Jari heavy cloud cover is common throughout the year, but we used the early and late time period imagery with the lowest cloud cover available to assess logging before and after logging activities occurred within the FMU (Table 1).

## 3. Methods

### 3.1 Data inputs for detecting selective logging

For the Landsat scenes given in Table 1, the surface reflectance values for the Blue, Green, Red, Near Infrared, Shortwave Infrared 1 and Shortwave Infrared 2 bands were measured at each pixel where logging occurred (*n*= 13699) and 2000 randomly selected pixels in an adjacent FMU that remained unlogged. In addition, since logging activities tend to be accompanied by surrounding disturbances (residual damage to neighbouring unharvested trees and skid trails along which logs are extracted), seven texture measures were calculated for each band (mean, variance, homogeneity, contrast, dissimilarity, entropy, and second moment) to provide a local context for each pixel (Beekhuizen and Clarke, 2010; Castillo-Santiago et al., 2010; Haralick et al., 1973; Rodriguez-Galiano et al., 2012). These were calculated within a 7×7 pixel window, chosen as a trade-off between minimizing window size while still capturing the disturbances in a selectively logged forest compared to an unlogged forest. The various texture metrics were assigned to the centre pixel, thus maintaining pixel size (i.e. 30 m), and were added after preliminary modelling efforts with only the surface reflectance bands were found to perform inadequately (i.e. approximately double the rate of omission error of logged pixels; see Table S1 for details). Because of possible Landsat inter-sensor differences, we added one final categorical variable that represented the sensor (TM, ETM+, or OLI) from which the image was acquired. The dataset thus comprised a 49-element vector (6 surface reflectance bands, 7 texture measures for each band, and a sensor-type indicator) for each pixel where logging occurred and an additional 2000 randomly selected pixels in an adjacent FMU that remained unlogged between 2008 and 2016.

The early and late datasets were reduced to exclude data from time periods close to when each FMU was logged. In the early dataset, for each FMU we excluded data from the year before logging because access roads were built and pixel values would therefore not represent undisturbed forest. In addition, data from all years following logging were excluded (see Table S2 for details). For example, for an FMU logged in 2014 the early dataset comprised data from around August in 2008 through 2012 (representative of unlogged conditions) and August 2014 (representative of logged conditions), but excluded data from August 2013, 2015 and 2016. The same procedure was used for the late dataset. For example, for an FMU logged in 2014 the unlogged dataset included data acquired around June in 2008 through 2012, while the logged data was for June 2015. Data were excluded from June 2013 (roads being built in the FMU), 2014 (logging recently initiated), and June 2016 (2 years post-logging). In both the early and late datasets the data from 2000 randomly selected pixels in an adjacent FMU that remained unlogged were retained from all years because they were never logged. Note that for early data, the imagery was acquired before the final part of the FMU was logged; this introduced some errors into model training, because some pixels labelled as logged in the training data were still unlogged. Despite this, we demonstrate in Section 4.1 that detection of selective logging was better with early time period data.

### 3.2 Random Forest for detection of selective logging

We built Random Forest (RF) models using the *randomForest* package in program R version 3.3.1 (Liaw and Wiener, 2002; R Development Core Team, 2016). The RF algorithm (Breiman, 2001) is a machine learning technique that uses an ensemble method to identify a response variable (here, whether a pixel was logged or unlogged) given a set of predictor variables (e.g. surface reflectance values). In contrast to a single decision tree, RF models employ multiple, independent decision trees (hence a forest). Random subsets of the training data are drawn, with replacement, to construct many trees in parallel, with each tree casting a vote on which class should be assigned to the input data. The withheld subset of the data, called the out-of-bag fraction, can be used for validation in the absence of independent validation data (Breiman, 2001). To reduce generalization error, RF also uses a random subset of predictor variables in the decision at each node within a tree during construction.

We split the early and late datasets into 75% for training and 25% was withheld for validation. We used the out-of-bag data during model training to determine the threshold value for classification (i.e. model calibration, see Section 3.3.1). In order to ensure independence, the training and validation datasets were spatially filtered such that no observations in the training dataset were within 90 m of an observation in the validation dataset. RF models have only two tuning parameters: the number of classification trees to be produced (*k*), and the number of predictor variables used at each node (*m*). We used 10-fold cross-validation to identify the number of trees (*k* = 1000) and the number of variables to use at each node (*m* = 5) that minimized the out-of-bag error rate on the training data.

### 3.3 Algorithm evaluation

#### 3.3.1 Calibration: selecting the detection threshold

RF models typically use a simple majority vote to assign an observation to a particular class, for example, in binary decisions when more than 50% of the trees assign a pixel to a particular class (Breiman, 2001). However, the proportion of votes cast for a particular class from the total set of trees can be obtained for each pixel and a classification threshold can be applied to this proportion (Liaw and Wiener, 2002). We adopted this approach here, wherein the proportion of votes that predicted each observation to be logged, denoted as *X* and informally termed the *likelihood* a pixel was logged, was used to select the classification threshold. Model calibration (with the out-of-bag data) was then used to define a threshold, *T*, such that if *X*> *T* the pixel was classified as logged (Fig. 3).

**Fig. 3.**
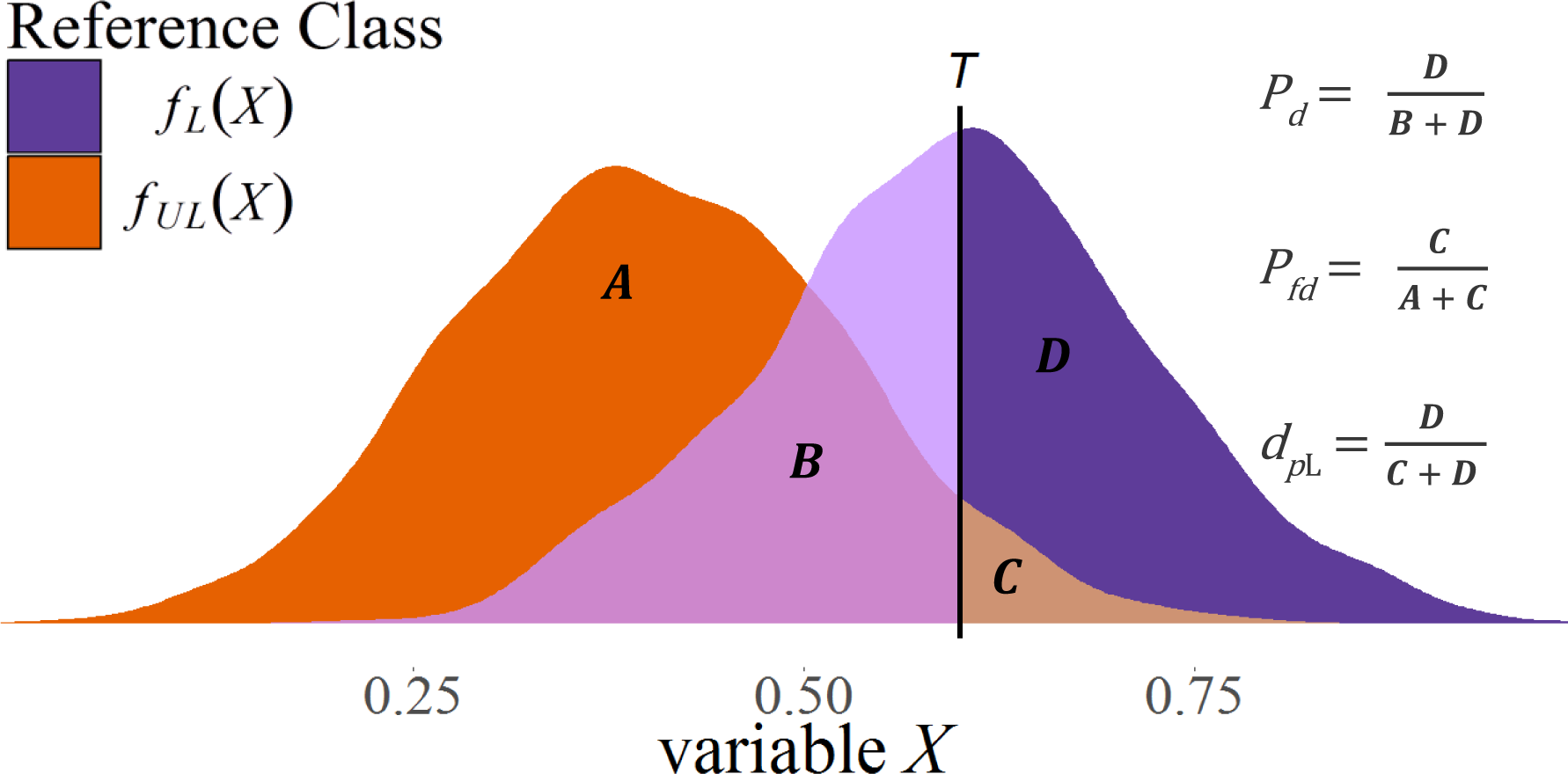
Diagram representing the trade-off between the probability of detection (*P_d_*) and the probability of false detection (*P_fd_*) associated with using a threshold *T* (vertical black line) on the variable *X* (the proportion of votes that predicted each observation to be logged) to label pixels as logged and unlogged. Here the purple and orange colors correspond to probability distribution functions of *X* for hypothetical logged, *f_L_*(*X*), and unlogged, *f_UL_*(*X*), observations, respectively. Thus, the areas A and B are the portions of the observations from unlogged and logged pixels, respectively, that will be labelled as unlogged. Similarly, C and D represent the portions of the observations from logged and unlogged pixels, respectively, that will be labelled as logged.

Detection of logging involves only two classes, logged and unlogged forest, so the confusion matrix has the form:

|  |  | Reference |  |
| --- | --- | --- | --- |
|  |  | L | UL |
| Predicted | L | $D_L$ | $D_{UL}$ |
| | UL | $N_L - D_L$ | $N_{UL} - D_{UL}$ |

where L and UL refer to logged and unlogged, *N_L_* and *N_UL_* are the numbers of logged and unlogged observations in the reference dataset, and *D_L_* and *D_UL_* are respectively the numbers of logged and unlogged pixels detected as logged. The total number of observations is *N* = *N_L_* + *N_UL_*. Since logging is a relatively rare event, both in our data and on the landscape (i.e. *N_L_* << *N_UL_*), it is appropriate to use the terminology of detection theory. Accordingly, we define the *detection probability P_fd_* = *D_L_/N_L_* and *false detection probability P_d_* = *D_UL_/N_UL_* as the probabilities that a logged or unlogged pixel is classified as logged, respectively. *P_d_* is equivalent to 1 – the omission error of the logged class and *P_fd_* is the omission error of the unlogged class.

A pixel was classified as logged if *X*, the proportion of votes from RF that predict the pixel as logged, exceeds a given threshold *T*. Hence the detection and false detection probabilities depend on *T* and can be written

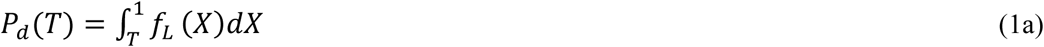

and

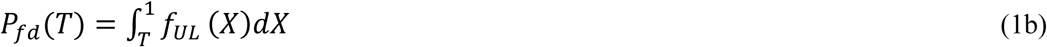

where *f_L_*(*X*) and *f_UL_*(*X*) are the probability distributions of *X* for the logged and unlogged classes, respectively (see Fig. 3).

The selection of *T* involves a trade-off between increasing *P_d_* and reducing *P_fd_* (Fig. 3). In making this choice, the overall accuracy, given by

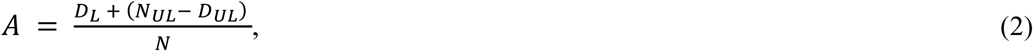

is not a good guide, since it can be shown that *A* is maximal (equivalently, the overall probability of error is a minimum) when

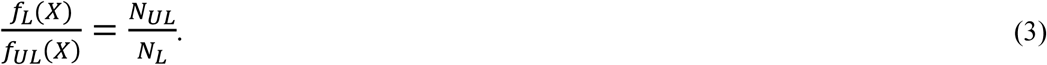

If *N_L_* and *N_UL_* were equal, the threshold would then be chosen at the intersection of *f_L_*(*X*) and *f_UL_*(*X*), but since *N_L_* << *N_UL_* it has a much higher value (i.e. it moves to the right in Fig. 3). This is because to increase overall accuracy it is more effective to reduce *P_fd_* than to increase *P_d_*, since there are so many more unlogged pixels (Schwartz, 1984), and maximizing accuracy would lead to very few (or even no) detections. For example, if only 1% of an area was logged and all the pixels were classified as unlogged, the overall accuracy would be 99%. Thus, overall accuracy would not sufficiently balance the trade-off between true and false detections to meet our objectives.

Various criteria could be used to select a classification threshold, including maximizing Cohen’s kappa (Cohen, 1960) or defining an acceptable rate of omission error; ultimately however, there is no wrong threshold, since this depends on the objectives of prediction. The criterion used in this study to define *T* was to fix the proportion of detected pixels that were truly logged, defined here as *d_pL_*:

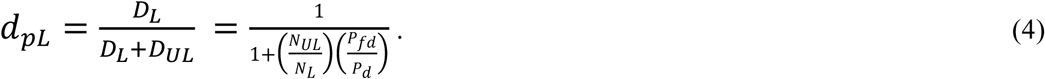

Adopting this criterion is equivalent to a Constant False Discovery Rate detector which is widely used in detection problems with rare events (Benjamini and Hochberg, 1995; Neuvial and Roquain, 2011). This fixes the rate of prediction error (i.e. type I) when labelling pixels as logged, because *d_pL_* is equal to 1 minus the commission error of the logged class, thus limiting the rate of commission error. This approach enables the user to select the proportion of detections that will be false. It was chosen because in the detection of rare events (e.g. selective logging within the Amazon Basin, for example), the implications of a particular error rate when predicting over the majority class (i.e. unlogged forest) are greater than an equivalent error rate when predicting over the minority class (i.e. 10% of millions of unlogged pixels is far greater than 10% of thousands of selectively logged pixels). Thus, in order to avoid being swamped by false detections, we wanted to fix the proportion of all detected pixels that were incorrect and accept the level of accuracy associated with this criterion. The approach outlined here, therefore, should be view from a detection theory perspective as opposed to simply being a classification problem.

Model calibration was used to calculate *P_d_, P_fd_*, and *d_pL_* across the full range of threshold values. In practice this involved iterating through all values of *T* between 0 and 1 (in steps of 0.001), building each confusion matrix, and calculating the associated values of *P_d_, P_fd_*, and *d_pL_*. The threshold value was chosen such that *d_pL_* = 0.85 in the training data (i.e. 15% of pixels classified as logged were actually unlogged). We initially set *d_pL_* to 95% to strongly limit the rate of false detections, but this resulted in very high omission error of truly logged pixels (>75%). Consequently, *d_pL_* was reduced to 0.85 by lowering the threshold, thus causing the detection and false detection rates to increase and causing more logged pixels to be detected. This value was then used to estimate *P_d_* and *P_fd_* during model assessment with the validation dataset.

#### 3.3.2 Validation: assessing model accuracy

RF models were validated using a random, independent subset of the early and late datasets (described in Section 3.2). The threshold value of *T*, chosen during model calibration, was applied to the validation data and the associated error rates were calculated. The values of *P_d_, P_fd_*, and *d_pL_* are presented across full range of threshold values to thoroughly illustrate model performance. Good practices outlined by Olofsson et al. (2014) were used to assess agreement and calculate unbiased error estimates when mapping selective logging detections. During mapping, non-forested areas were excluded using Brazil’s national forest change product, PRODES (INPE 2015), and cloudy pixels were masked using the cloud mask provided with Landsat surface reflectance imagery. In addition, we provide the value of Cohen’s kappa, κ, for comparison with other studies (Cohen, 1960).

### 4. Results

#### 4.1 Random Forest classification of selective logging at Jamari

The rates of true and false detection probabilities for the early and late validation data are shown in Fig. 4 for the full range of *T* (black lines). These curves indicate how a given threshold value used for classification influenced the associated values of *P_d_, P_fd_*, *κ*, and *d_pL_* in the validation data. For example, if a *d_pL_* of 0.90 was used (indicating 10% of logging detections would be spurious) then the false detection rate (*P_fd_*) would be < 1% for both datasets, but the detection rate (*P_d_*) would be approximately 55% and 30% for the early and late datasets, respectively. These plots clearly demonstrate that there is no unambiguous way to choose an optimal value for *T*, and the choice about its value is a trade-off between the number of true and false detections.

**Fig. 4.**
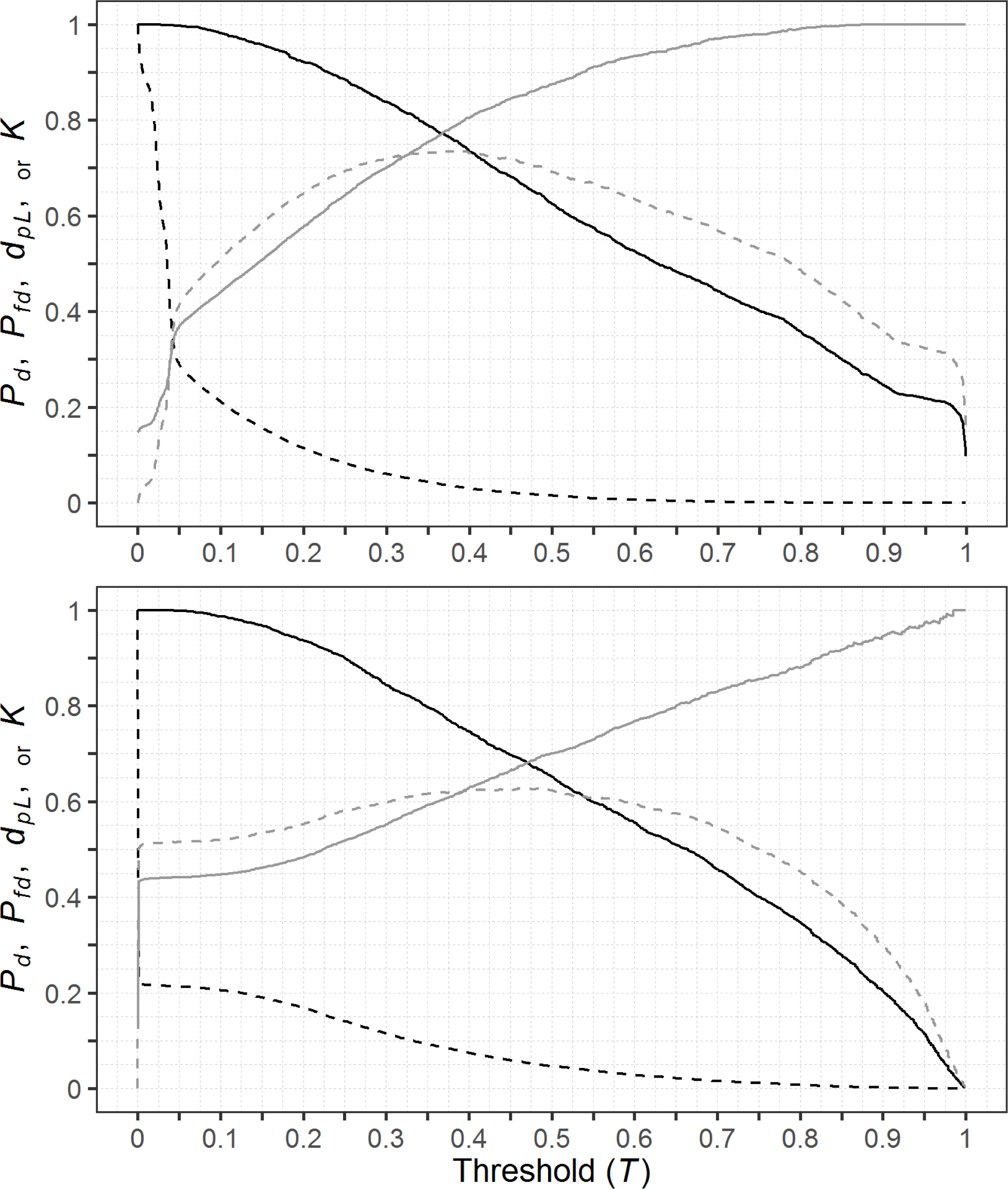
Trade-off curves between true (*P_d_*) and false (*P_fd_*) detection rates (solid and dashed black lines, respectively) for the early (top) and late (bottom) Random Forest models at the Jamari site as a result of varying the threshold value (*T*) for classification. Also shown are the corresponding values of *d_pL_* (the proportion of detections that were truly logged) and Cohen’s kappa (solid and dashed grey lines, respectively).

In general, these plots indicate that the early data provided a higher detection rate than the late data, for a given false detection rate. The early and late data had similar rates of commission error when labelling logged pixels, which is not surprising given we used this measure to constrain models during training. However, the late data had higher rates of omission error of logged pixels and detected less logging (Table 2). In addition, these plots demonstrate why using the threshold that maximized Cohen’s κ would lead to higher false detection rates, as the threshold value is higher when *d_pL_* = 0.85 than at maximum κ (i.e. pixels classified as logged must have a higher likelihood). Furthermore, because κ is high across a wide range of range of threshold values for both early and late data, slight differences in the likelihoods produced by the validation data could result in dramatic shifts in the value of *T*.

Although *d_pL_* was fixed at 0.85 during model calibration (i.e. with the training data), the values calculated with the validation dataset were slightly lower (Table 2). Thus, the threshold value determined during model training did not produce the same values for *d_pL_* when used against the validation dataset (i.e. some loss of performance). Slight differences in the proportion of logged observations (16.3% and 14.5% in training and validation, respectively) and minor differences in the ratio of *P_fd_*: *P_d_* between the training and validation datasets account for the disparity (see equation 4). In general model assessments seldom give identical performance across training and validation phases, and the difference here were marginal and yielded comparable model behavior.

The early data displayed higher spatial correspondence between high likelihoods and the locations of logging in Jamari. This is illustrated in Fig. 5, where the likelihood of logging provided by RF is shown on a colour scale and the individual locations of tree removal are indicated by black squares. The early model yields much higher likelihoods and these match well with reference logging data, whereas there is generally lower correspondence between reference logging locations and regions of highest likelihood in the late predictions. Note that we expect some logging locations to be omitted in the early data as the corresponding satellite data were acquired part way through the logging period and missed later logging. Evidence for this is provided by the inset regions expanded at the bottom of Fig. 5 where the locations of the last 200 trees in the logging records for the season are displayed as crosses instead of squares. Many of these locations occur in low likelihood regions in the early data because these locations were probably unlogged at the time of the image acquisition (dates for specific tree removal were unavailable).

**Fig. 5.**
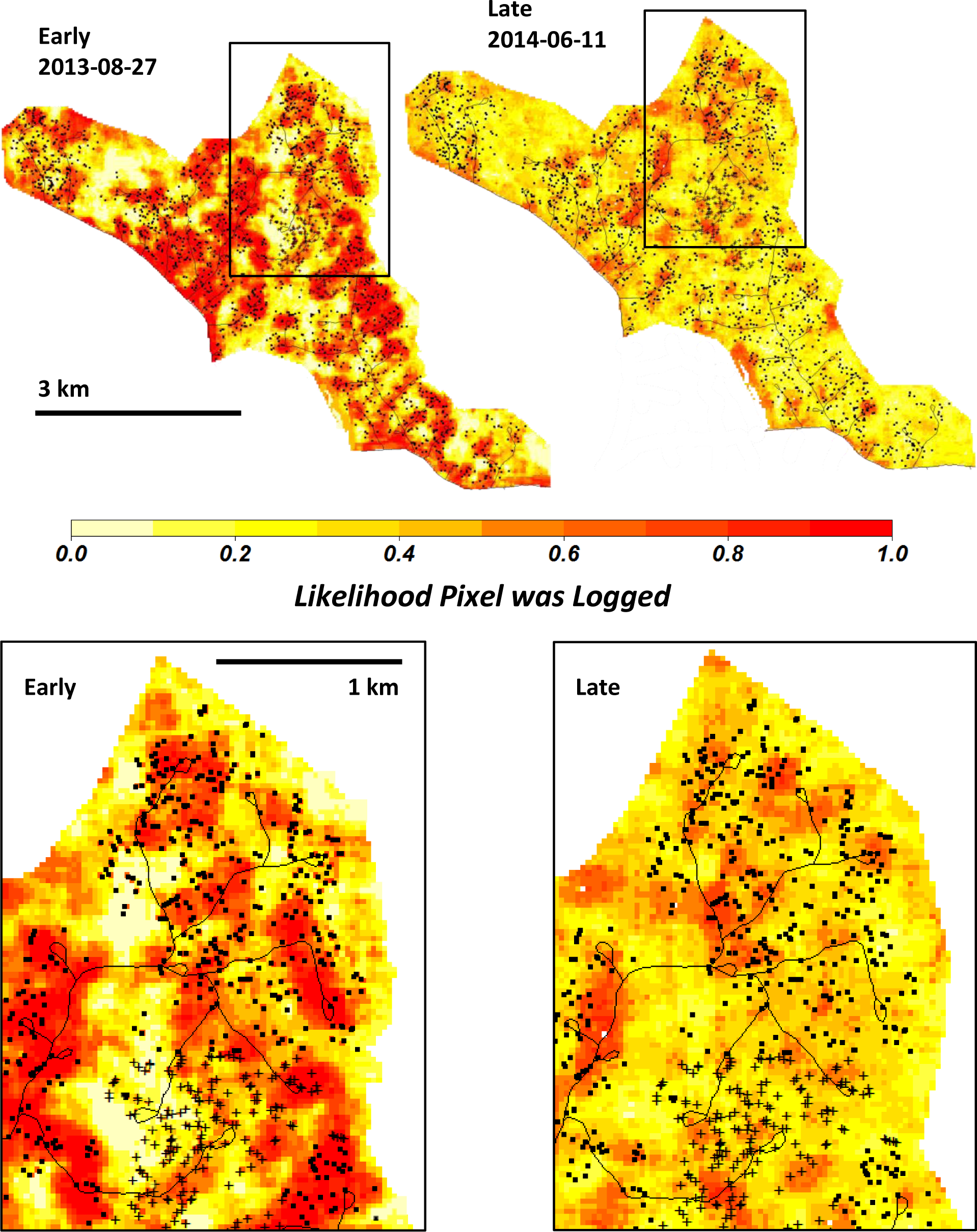
Example of a forest management unit in Jamari logged in 2013 showing the RF predicted likelihood that each pixel was logged (highest likelihoods in red) for the early and late data. Logging roads are thin black lines and tree removal locations are displayed as black squares and crosses. The black crosses (see insets for detail) coincide with the final 200 trees in the logging records for 2013.

**Table 2.** Confusion matrix summarizing unbiased (Olofsson et al., 2014) results from Random Forest (RF) model classifications of logged and unlogged observations at Jamari derived from Landsat data at labelled points (observations before and after selective logging). The classification threshold (*T*) for RF models was set during model calibration such that the proportion of detections that were truly logged (*d_pL_*) was fixed at 0.85, resulting in a *T* of 0.40 and 0.65 for the early and late datasets, respectively. The corresponding values for overall accuracy (OA), Cohen’s kappa (κ), the proportion of detected pixels that were truly logged (*d_pL_*), and the detection probability (*P_d_*) are provided.

| EARLY |  |  |  |  | LATE |  |  |  |  |
| --- | --- | --- | --- | --- | --- | --- | --- | --- | --- |
| OA: 89.7% |  |  |  |  | OA: 91.7% |  |  |  |  |
| $\kappa$: 0.78 | | | | | $\kappa$: 0.40 | | | | |
| $d_{pL}$: 0.80 | | | | | $d_{pL}$: 0.80 | | | | |
| $P_d$: 0.92 | | | | | $P_d$: 0.30 | | | | |
|  |  | Reference Class |  | Commission Error (%) |  |  | Reference Class |  | Commission Error (%) |
|  |  | Logged | Unlogged |  |  |  | Logged | Unlogged |  |
| Predicted | Logged | 0.313 | 0.076 | 19.5 | Predicted | Logged | 0.032 | 0.008 | 19.9 |
| Class | Unlogged | 0.027 | 0.584 | 4.4 | Class | Unlogged | 0.075 | 0.885 | 7.8 |
| Omission Error (%) |  | 8.0 | 11.5 |  | Omission Error (%) |  | 70.1 | 1.0 |  |

A further marked difference between the predictions is that, in general, far more pixels were labelled as logged in the early data than in the late, as can be seen by comparing the classifications in Fig. 6, which shows the years between 2011 and 2015 for the early (top) and late (bottom) datasets, respectively. The FMU where logging occurred in each year is outlined in yellow and the 2015 image also shows the FMU to be logged in 2016 outlined in white. The early classifications appear to show some indication of a retained signal from the previously logged FMU (particularly 2012-08-16 and 2013-08-27 in Fig. 6) that are less visible in the late classifications. In addition, the range of predicted logging likelihoods with late data was more variable from scene to scene, which resulted in some scenes having very few pixels of high likelihood of logging (see 2012-06-13 in Fig. 6) and others with most of the study area predicted as logged (see 2016-06-16 in Fig. 6). This suggests the threshold value from model calibration could not be used reliably for all late images and a scene-specific threshold value might need to be calculated for each image to provide better correspondence with logging activities.

**Fig. 6.**
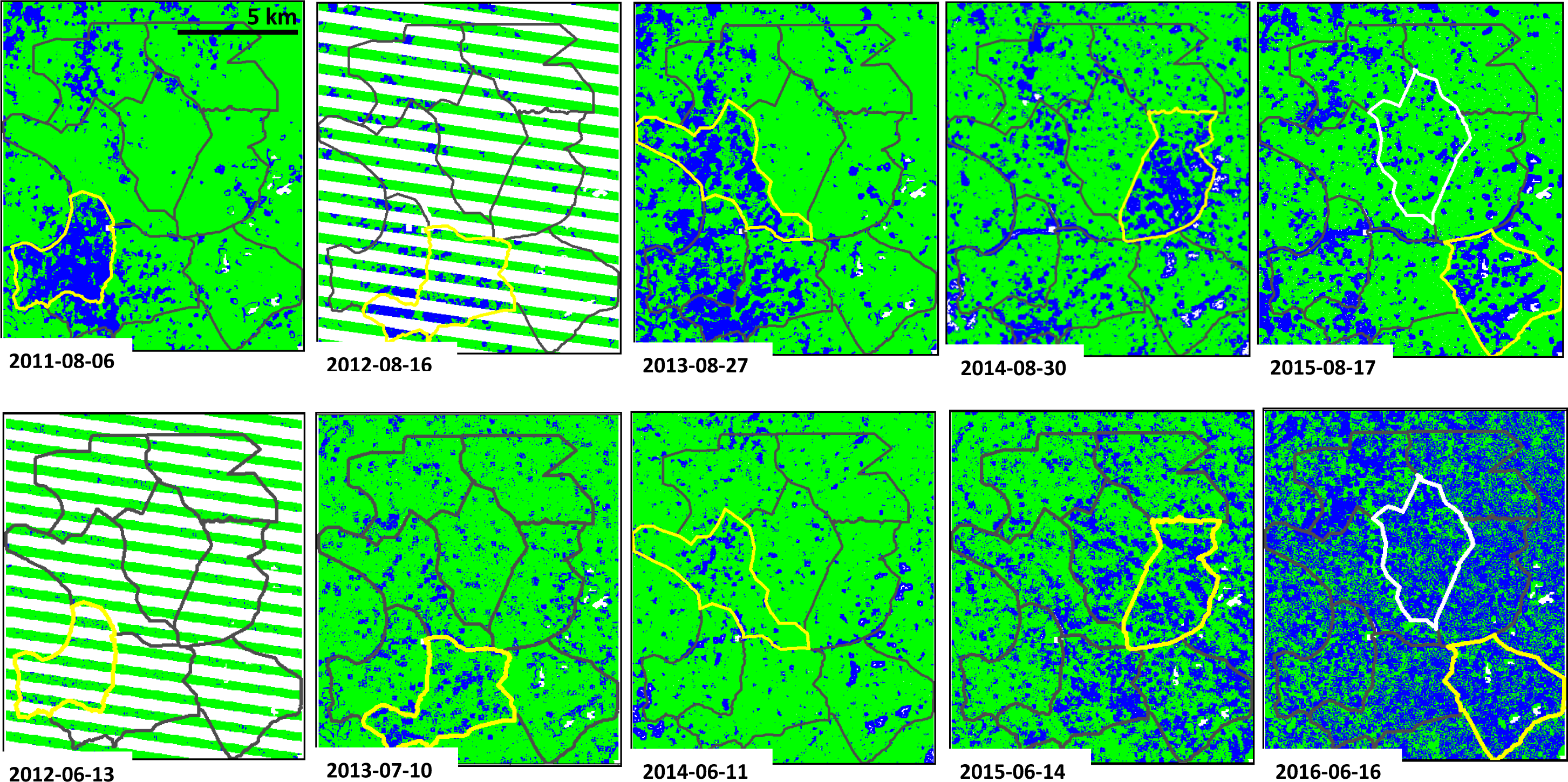
Classifications for Jamari between 2011 and 2016 with early (top) and late (bottom) Landsat data. The forest management units (FMUs) are outlined in black and the FMU logged in each year (where logging should be detected) is outlined in yellow. Blue and green represent classifications for logged and unlogged forest, respectively. White areas are no-date and correspond to the Landsat 7 scan-line corrector error (stripes) and pixels that were non-forest (irregular patches) in Brazil’s Program to Calculate Deforestation in the Amazon (PRODES) database. The FMU logged in 2016 is outlined in white (far right) and the top two FMUs in each image remained unlogged.

The true proportion of logged pixels in each FMU (from the logging records) was roughly 12% in a given year (mean = 11.8%; standard deviation = 2.4%), but the early classifications consistently labelled a greater number of pixels as logged (Fig. 7). For example, the proportion of pixels assigned in each FMU for early acquisitions was expected to be around 25% (10% truly logged and 15% false positives), but nearly twice as many were identified. However, forest disturbances from selective logging affect patches of forest and not just the pixels where trees were logged. Extra detections would be expected because of additional tree and canopy damage associated with tree removals, roads, and construction of skid trails. Note that the rate of false detections over unlogged FMUs (open diamonds in Fig. 7) is roughly as expected for the early algorithm and most dates for the late algorithm, but is significantly different for the late algorithm for the FMU logged in 2015. The late scene for this FMU clearly shows anomalous behaviour and displays high likelihood of logging over most of the study area, including known unlogged regions (see Fig. 6).

**Fig. 7.**
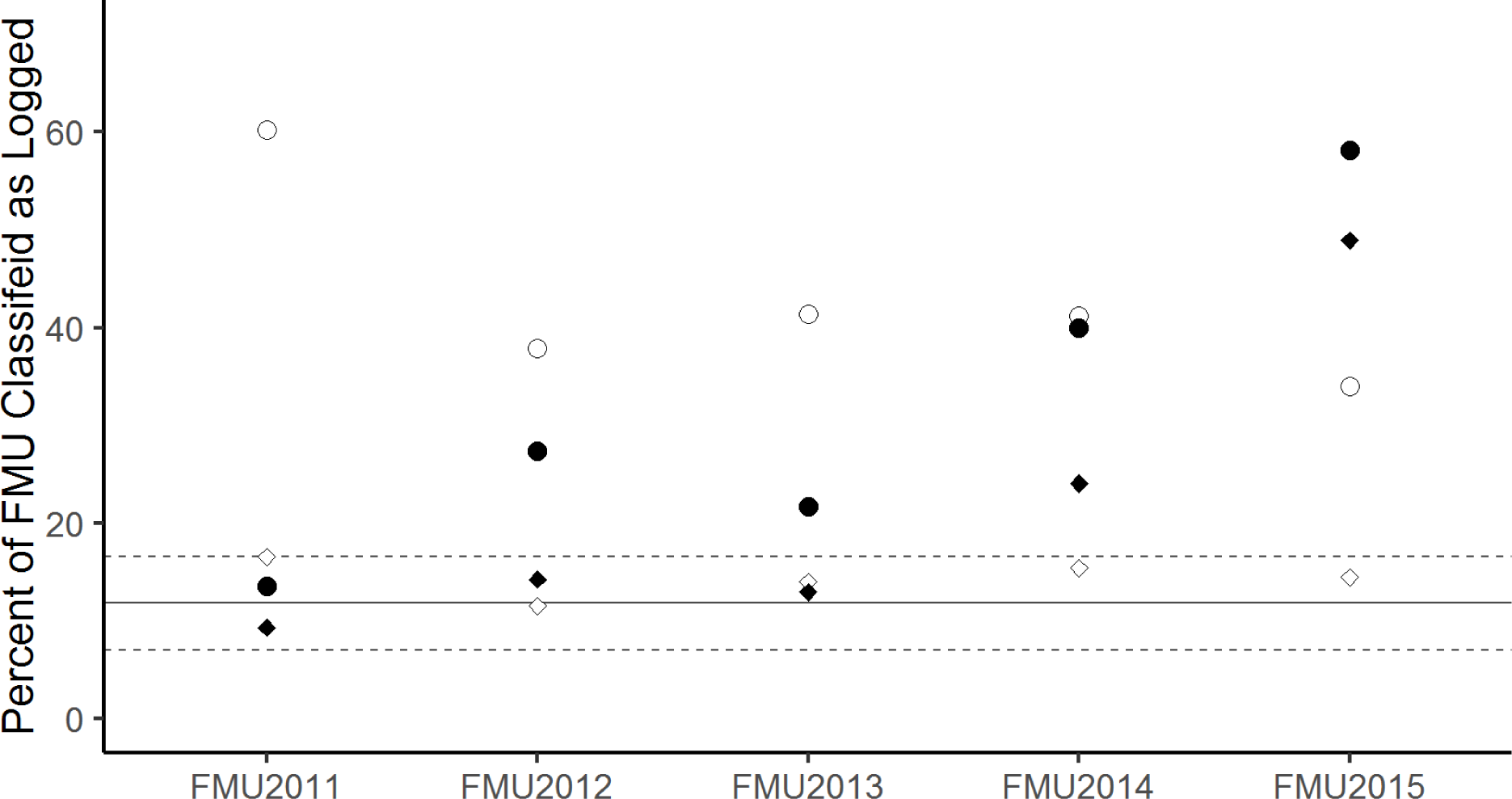
The proportion of pixels in each FMU that were classified as logged in Fig. 6 for the early (open symbols) and late (closed symbols) algorithms. Circles are the logged FMUs in each year and diamonds are values from an FMU that remained unlogged. The black line represents the mean ± 1 standard deviation (dashed lines) of the true rate of logging across all FMUs. Values are unbiased (Olofsson et al., 2014) to account for possible sampling bias in the validation data.

We used the early algorithm to predict over the available Landsat time series in Jamari that coincided with logging in four FMUs (see Table S3 for image dates) and plotted the detections of logging through time (Fig. 8). As expected, the proportion of detected pixels increased through the logging season during the year a given FMU was logged. There was also a drift upwards in the unlogged FMU, but the detections peaked just above the expected rate of 12% by late August (Fig. 8). Importantly, known unlogged regions will not exhibit a *d_pL_* of 0.85 (i.e. a false discovery rate of 15%), as any and all detections in known unlogged areas are wrong (i.e. a *d_pL_* = 0). Consequently, the false alarm rate is the expected proportion of detections (i.e. *P_fd_* = 11.5% in Table 2). This suggests that the algorithm performed as would be expected for tracking forest disturbances through time in both logged and unlogged FMUs. In particular, forest patches subjected to selective logging should display measurable increases in detections as the logging season progresses and known unlogged regions will exhibit the expected false alarm rate.

**Fig. 8.**
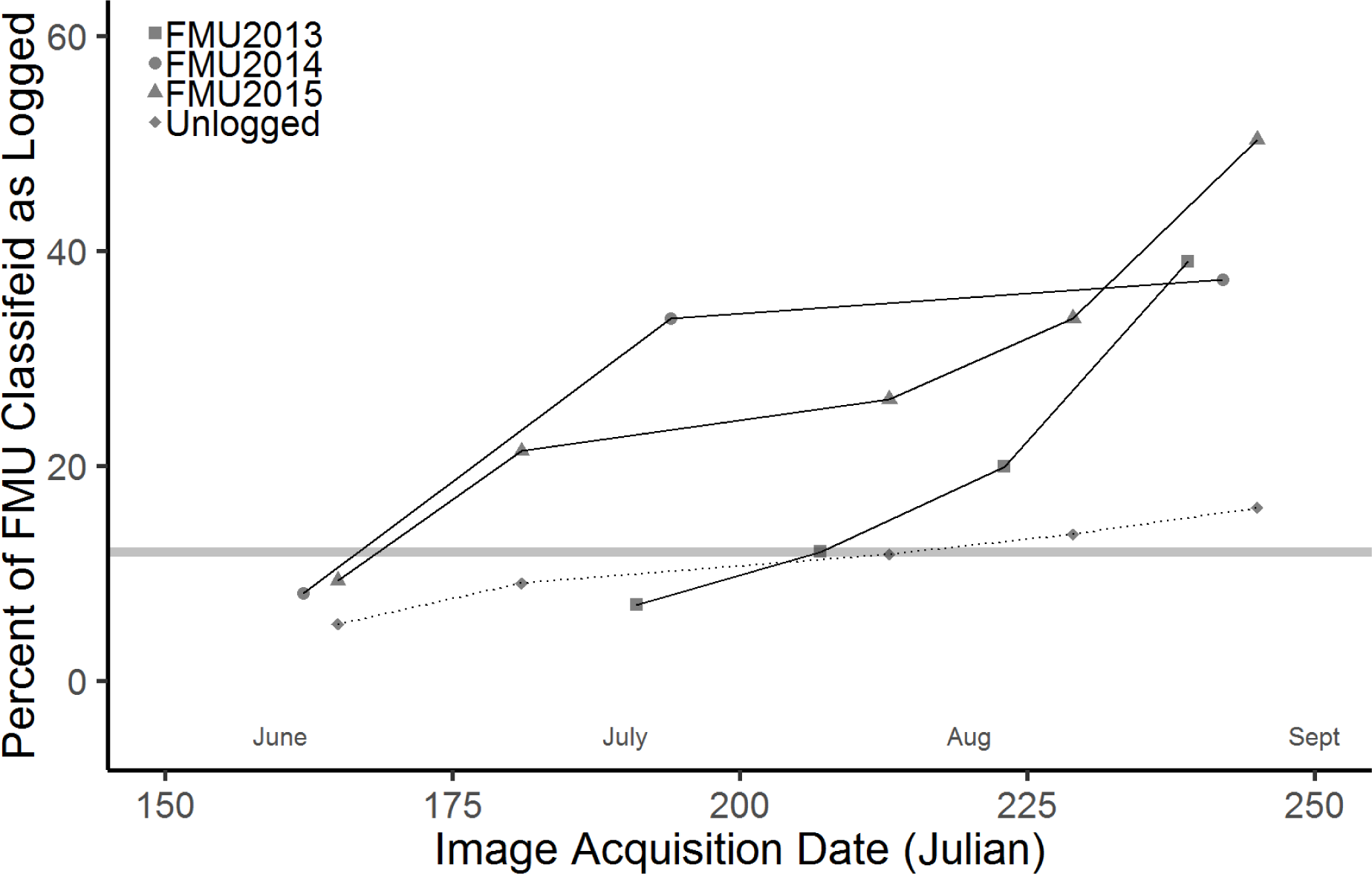
The proportion of pixels classified as logged through time in three logged and one unlogged FMU using the early RF model. Triangles, circles, and squares represent logged FMUs (solid lines) and diamonds are an unlogged FMU (dotted line). The grey horizontal line at 12% is the approximate detection rate expected for unlogged regions. Values are unbiased (Olofsson et al., 2014) to account for possible sampling bias in the validation data.

We assessed the impact of the window size used to calculate texture measures on the proportion of pixels labelled as logged FMUs for three logged and one unlogged FMU in the early data (Fig. 9). Reducing the window size from 7×7 to 3×3 lowered the proportion labelled as logged by nearly 50% within each FMU, resulting in smaller clusters of pixels with high likelihoods (Fig. 9). However, as noted above, forest disturbance from selective logging affects chunks of forest and not just the pixels where trees are cut. Thus, depending on the scale of interest, larger or smaller window sizes may be better for identifying patches of forest that have been selectively logged. In contrast, reducing the window size had little impact on the false detection rate over unlogged regions, remaining close to the 12% expected irrespective of window size (Fig. 9). This suggests that the choice of window size is independent of the false positive rate over undisturbed forested areas and primarily affects likelihoods around pixels that the algorithm identifies as disturbed.

**Fig. 9.**
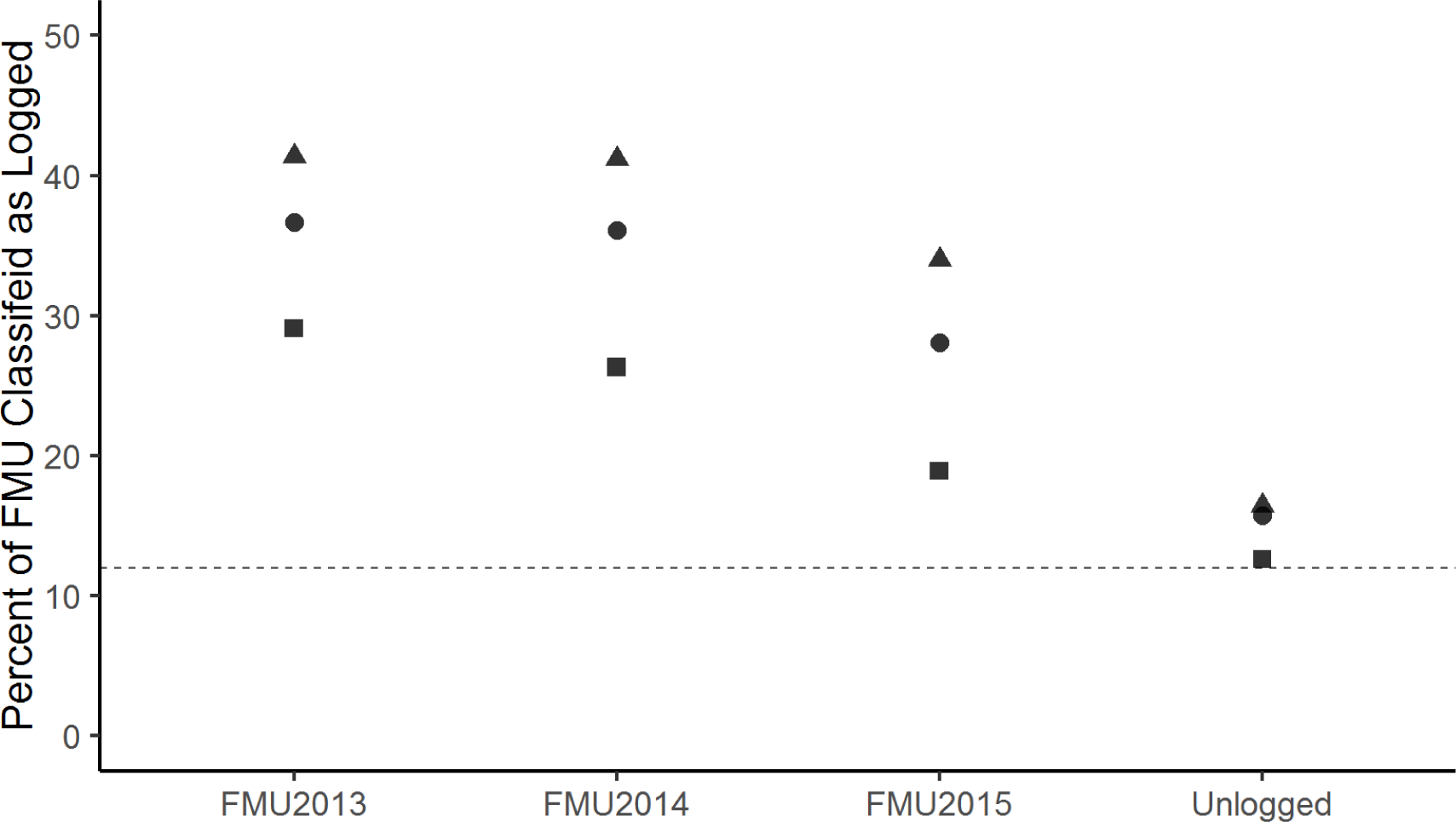
The proportion of pixels classified as logged in three logged FMUs and one unlogged FMU from RF models using texture measures with different window sizes. Triangles, circles, and squares represent windows used for texture calculation of 7×7, 5×5 and 3×3 pixels, respectively. The dashed line at 12% is the approximate detection rate expected for unlogged regions. Values are unbiased (Olofsson et al., 2014) to account for possible sampling bias in the validation data.

#### 4.3 Random Forest predictions of logging at Jari

The majority of the best available (sufficiently cloud-free) Landsat scenes over Jari were from the ETM+ sensor, which suffered the scan-line corrector error, so approximately 22% of each image has missing data that appear as white stripes in Figs. 10 and 11 (Storey et al., 2005). Nonetheless, this allowed us to see behaviour similar to Jamari, wherein predictions using early data clearly identified active logging (Fig. 10) and predictions using late data detected very little logging (Fig. 11). In particular, with late data most of the study area was classified as unlogged both before and after logging. Additionally, with early data the predictions of logged pixels in the year before logging were close to the expected rate of false positives over unlogged regions (approximately 12%). However, with late data the rate of false positives was not close to the expected rate over unlogged regions. Maps for the year before logging are displayed to demonstrate that the early dataset identified the correct year in which logging occurred and did not simply predict high amounts of logging for every year.

**Fig. 10.**
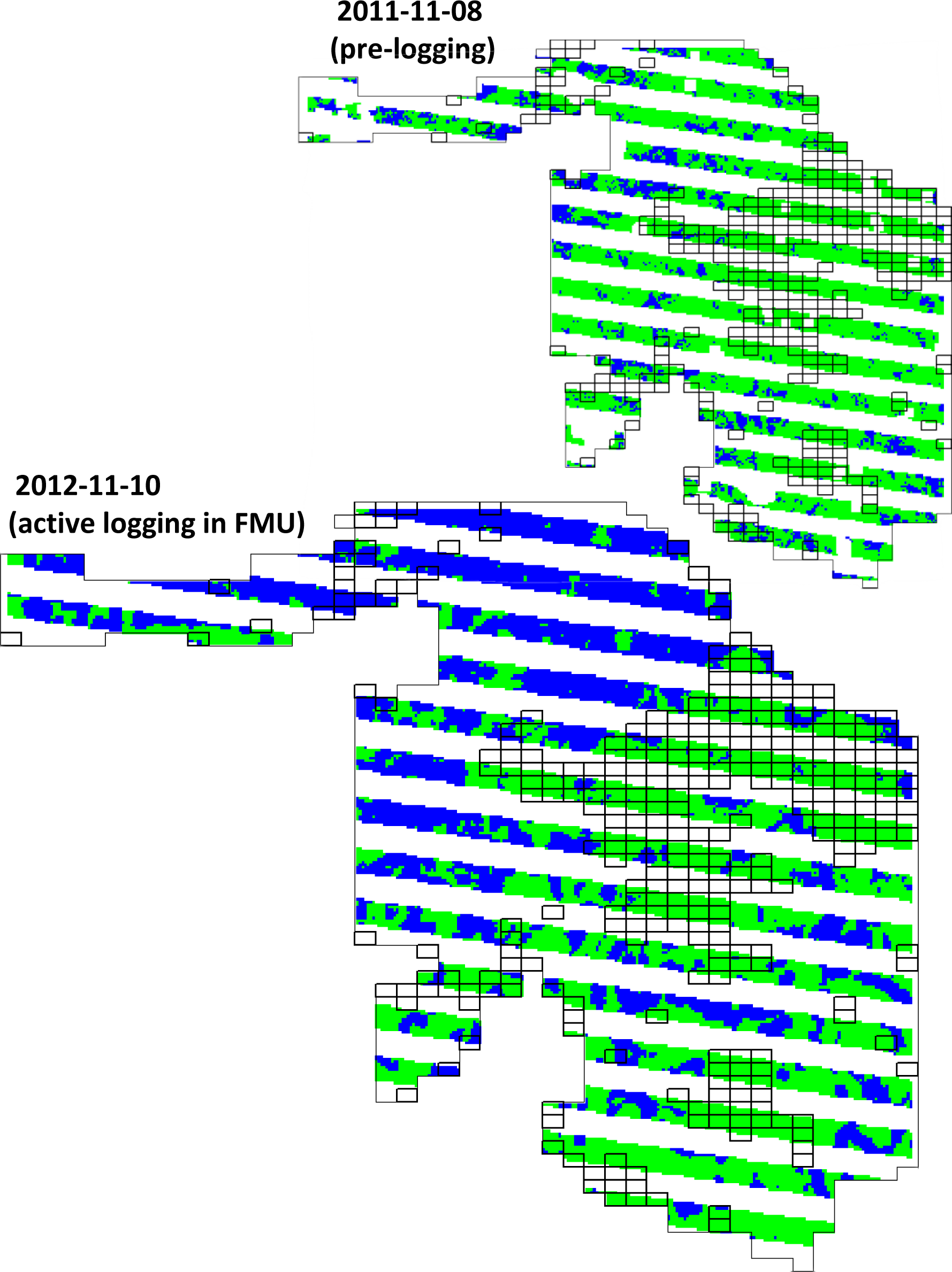
Logged (blue) and unlogged (green) predictions at the Jari study site using a Random Forest model trained from early Landsat inputs. Predictions from November 2011 (top) were before logging activities began and from November 2012 (bottom) while active logging was ongoing. Clouds were masked out and appear as irregular white patches (top). Missing data regions from the Landsat 7 scan-line corrector error appear as white stripes through the maps. Black boxes indicate the 10 ha blocks inside the Jari concession that were not logged.

**Fig. 11.**
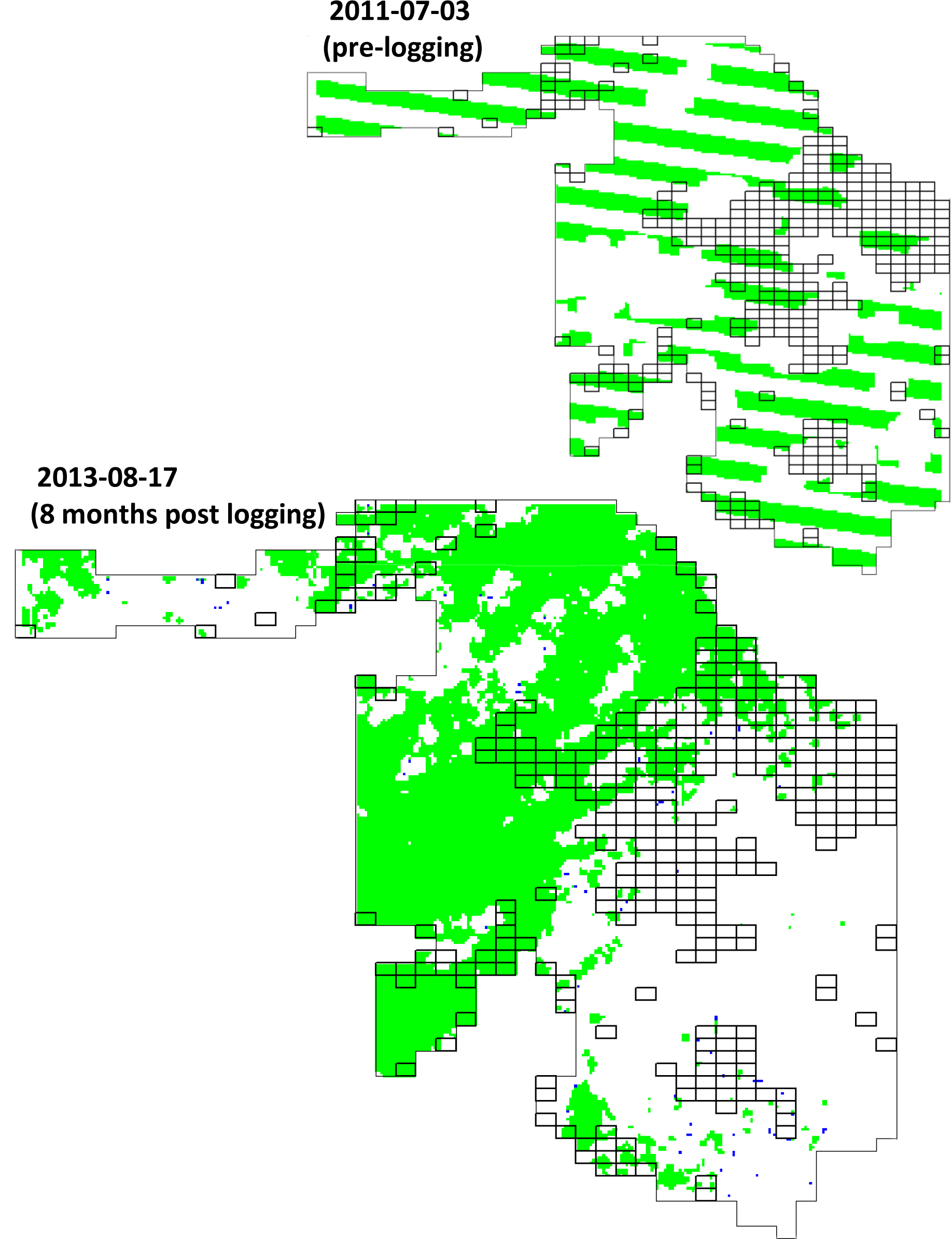
Logged (blue) and unlogged (green) predictions at the Jari study site using a Random Forest model trained from late Landsat inputs. Predictions from July 2011 (top) were before logging activities began and from August 2013 (bottom), approximately 8 months post-logging. Clouds were masked out and appear as irregular white patches. Missing data regions from the Landsat 7 scan-line corrector error appear as white stripes through the map (top). Black boxes indicate the 10 ha blocks inside the Jari concession that were not logged.

In total, an area of 6152 ha was visible in Jari after removing clouds and missing data gaps from the SLC error in the year of logging. Of this area, 1710 ha was not logged (black boxes in Figs. 10 and 11). Since we lacked detailed logging records and only knew which 10 ha blocks were logged, a formal accuracy assessment of logging detections was not possible. However, when using the unbiased proportions and the threshold from Table 4 to classify predictions, the early algorithm labelled 2316 ha (38%) as logged (Fig. 10). This value is consistent with predictions from Jamari where approximately 40% of logged FMUs were labelled with early data (see Fig. 7). In addition, the rate of commission error when predicting logged pixels (i.e. 1 - *d*_*p*L_) was 19.8%, which is also consistent with the rate of commission error between the validation data and prediction errors found for the Jamari site.

**Table 3.** Confusion matrix summarizing unbiased (Olofsson et al., 2014) results from Random Forest (RF) model classifications of logged and unlogged observations at Jari with Landsat data. The thresholds (*T*) developed at Jamari were used to classify predictions at Jari and were 0.40 and 0.65 for the early and late datasets, respectively (Table 2). The corresponding values for overall accuracy (OA), Cohen’s kappa (κ), the proportion of detected pixels that were truly logged (*d_pL_*), and the detection probability (*P_d_*) are provided.

| EARLY |  |  |  |  | LATE |  |  |  |  |
| --- | --- | --- | --- | --- | --- | --- | --- | --- | --- |
| OA: 89.0% |  |  |  |  | OA: 92.2% |  |  |  |  |
| $\kappa$: 0.77 | | | | | $\kappa$: 0.05 | | | | |
| $d_{pL}$: 0.80 | | | | | $d_{pL}$: 0.80 | | | | |
| $P_d$: 0.93 | | | | | $P_d$: 0.03 | | | | |
|  |  | Reference Class |  | Commission Error (%) |  |  | Reference Class |  | Commission Error (%) |
|  |  | Logged | Unlogged |  |  |  | Logged | Unlogged |  |
| Predicted Class | Logged | 0.351 | 0.085 | 19.5 | Predicted Class | Logged | 0.002 | 0.005 | 19.9 |
|  | Unlogged | 0.025 | 0.538 | 4.4 |  | Unlogged | 0.078 | 0.919 | 7.8 |
| Omission Error (%) |  | 6.7 | 13.7 |  | Omission Error (%) |  | 97.3 | 0.06 |  |

### 5. Discussion

The spatial resolution of Landsat data has previously been considered too coarse to monitor selective logging activities (Asner et al., 2002), with most applications involving logging intensities >20 m^3^ ha^−1^ at sites with an abundance of spectrally distinct features (Asner et al., 2005; Souza and Barreto, 2000; Souza et al., 2005). However, we have demonstrated that Landsat surface reflectance data can be used effectively, in a supervised machine learning framework, to detect subtle spectral changes from selective logging at low intensities. Although a definitive estimate of the amount of logging activities that have previously gone undetected is difficult to determine, a dataset of 824 logging permits from the state of Pará, Brazil found 18% of permits authorized for logging were harvested at intensities < 20 m^3^ ha^−1^ (Richardson and Peres, 2016). Thus, our approach has the potential to significantly increase current abilities to detect and monitor selective logging activities that up to now have been, at best, marginally detectable (see Supplementary Material, Section 3 for a comparison between our method and CLASlite, Asner et al., 2009a). In addition, the approach outlined here has the distinct advantage of being able to make predictions about forests on a single scene to map disturbances, instead of requiring successive cloud-free images like many approaches (Asner et al., 2009a). This is particularly important since a single, low-cloud scene may be all that is available for a given region (see Souza, Jr et al. 2013).

Only the algorithm developed with data close to the time of active logging (i.e. the early data) performed well at detecting selective logging. Many logged pixels were omitted when using data from the first cloud-free image of the next dry season (i.e. late). In addition, only the algorithm trained with imagery close in time to the logging events was transferable to new areas (Figs. 10 and 11). Thus, our results suggest images acquired during, or very soon after, active logging are needed to map low intensity selective logging. This is partly because logging activities typically occur in the dry season when cloud-free imagery is more likely to be available, but also because the spectral changes associated with low-intensity selective logging practices are subtle and short-lived and rapidly become obscured under even limited regrowth (Broadbent et al., 2006).

The decision to fix the proportion of logging detections that were correct (i.e. limiting the commission error when predicting logged pixels) defined the classification threshold applied to the likelihoods produced by the RF models developed at Jamari. This threshold would likely give different values of *d_pL_* in regions that contain different proportions of logged and unlogged observations (see equation 4). Indeed, the threshold value from model training produced a slightly higher *d_pL_* when assessed against the validation dataset, yet these data were from the same study site. In addition, depending on the distribution of likelihoods produced by the RF models, different datasets might yield different threshold values, for example because of higher selective logging intensities. However, assuming both classes are present, the proportion of detected pixels that are wrong (i.e. 1 - *d_pL_*) would be expected to remain invariant. Hence if the same threshold were applied over the whole of the Amazon basin, we would expect approximately 20% of all detections to be wrong and 11.5% of truly intact forest pixels to be identified as logged. This could be used to refine the algorithm (in the absence of field data on logging locations) by examining the rate of false detections over known unlogged regions or protected areas to achieve a similar error rate. Adopting this threshold (i.e. *P_fd_* = 11.5) would make the method equivalent to a Constant False Alarm Rate detector which is widely used in detection problems with rare events (Scharf, 1991). A *d_pL_* of 85% was the value chosen here as a compromise that gives a high detection rate (0.92 for early data, see Table 2) while keeping the proportion of detections that are false to an acceptable level. However, other values of *d_pL_* could be chosen, depending on the predictive objectives of the particular application. This is precisely why Fig. 4 shows the full range of threshold values; to enable a detailed assessment of model performance with higher or lower values of *T* or *d_pL_*.

An important issue when assessing detections of selective logging is that *patches* of forest are affected, not just the isolated pixels where trees are removed. The area around logged pixels is certain to be disturbed because of canopy damage associated with tree removals and the construction of roads and skid trails, but the precise amount is unknown. Consequently, taking as a reference purely the pixels where trees were known to be removed is inadequate for assessing the disturbance due to logging. Indeed, the true rate of logged pixels at Jamari was approximately 12% (mean = 11.8%; standard deviation = 2.4%), but this represents a minimum expected detection rate and the associated forest disturbances would result in more detections. The early algorithm labelled approximately 40% of the area inside FMUs in Jamari and Jari as logged. This may be a more realistic estimate and is likely close to the upper limit of what constitutes forest disturbance for this level of logging. However, because the choice of window size for texture measure calculation affected the proportion of pixels labelled as logged (Fig 9), the appropriate window size for a particular application needs to be considered. Smaller windows resulted in fewer detections, but use of too small a window risks being unable to adequately measure texture arising from forest disturbances from selective logging. Thus, the specific application would best dictate the optimum approach and the user should, if possible, use window sizes matched to the expected or known spatial spread of forest disturbance around tree removals.

Selective logging rates in the Brazilian Legal Amazon (BLA) are thought to have remained relatively stable since 2000, with Pará and Mato Grosso enduring the highest rates of selective logging (Betts et al., 2017; Souza, Jr et al., 2013). However, our findings suggest that their assessments of forest disturbance and the associated carbon emissions are likely underestimated. Machine learning approaches (neural networks, decision trees, support vector machines, etc.) for classification of satellite imagery have been used with increasing frequency and success since their initial applications to remote sensing questions in the 1990’s (Tuia et al., 2011), but their effectiveness relies heavily on adequate training data. Our results suggest that detailed logging records ought to be a reporting requirement for logging companies or for REDD+ projects related to logging. These datasets could be used for building, improving, and updating models similar to the one presented here, with the aim of facilitating the creation of pan-tropical estimates of (legal and illegal) selective logging activities.

From a conservation perspective, the ability to identify regions of forest that are selectively logged is useful for mapping primary forest, but also for delineating logged forests with conservation value. Forests subjected to selective logging generally maintain far higher levels of biodiversity than other modified habitats, such as plantations or secondary forests (Gibson et al. 2011; Edwards *et al.*, 2014). Moreover, even after accounting for the amount of wood removed, reduced impact logging activities (like those at our study site in Jamari) do better at maintaining biodiversity than conventional selective logging practices (Bicknell et al., 2014) while simultaneously sequestering more carbon during regrowth (Martin et al., 2015; Putz et al., 2008). Thus, in the context of REDD+ or alternative conservation initiatives, forests affected by low intensity selective logging offer high biodiversity value and carbon sequestration potential. Accordingly, our method could be used for identifying and prioritizing forest tracts suitable for such initiatives.

#### 5.1 Study limitations

While the minimum mapping unit remained 30 m, the use of texture measures resulted in some spatial aggregation of logging predictions (see Figs. 5 and 6). This was expected around logged pixels, as a result of canopy gaps, skid trails, and roads, but clustered detections were also present in unlogged FMUs (see Fig. 6). Ideally, predictions of logging in unlogged FMUs would have shown a diffuse 12-15% of spurious detections. Attempts to refine the accuracy of a final predictive map, by performing a post-processing step in which either likelihoods or classified pixels are re-examined (e.g. using a window analysis to apply neighbourhood rules whereby likelihoods or counts of nearby pixels are reevaluated against some criteria) to enhance the detection rate or limit the false detection rate further, would prove difficult (Huang et al., 2014). However, using a smaller window size for texture calculation, such as 5×5 pixels, would reduce this effect. Ultimately, the optimal window size for textures depends on the objectives of the application and understanding how different window sizes affect detection and false detection rates.

Landsat surface reflectance data is known to exhibit occasional strong scene-to-scene and within-scene variations because of discontinuities across focal plane modules (Morfitt et al., 2015) and seasonal changes in solar viewing angles (Roy et al., 2016), respectively. We did not take these effects into account and likely affected algorithm performance in some instances (e.g. 2016-06-16 in Fig. 6). Thus, a large scale application of the approach outlined here should include a step to normalize surface reflectance data across scenes to facilitate detection of the subtle and short lived spectral changes associated with low-intensity selective logging practices (Broadbent et al., 2006).

Our analysis used a binary classification (logged and unlogged forest) yet tropical forest landscapes are a heterogeneous mixture of land uses (e.g. secondary forests, burned areas inside forests, agricultural fields). We avoided some of these complexities by using the PRODES forest designations to remove urban areas, agricultural fields, and deforested areas that had regenerated to secondary forest. However, our method cannot distinguish between disturbance types and is best suited for tracts of remaining forest that contain logging concessions. In addition, selective logging represents a range of forest disturbance intensities and we would have preferred to use the logging dataset in a regression framework (i.e. a continuous response, such as logging intensity). However, the range of logging intensities within our Jamari dataset was very limited, since it was such a low intensity concession. Consequently, a regression approach was not suitable for the Jamari dataset and we chose to use classification. Additional datasets could fold into the framework here and might facilitate a continuous response approach as those datasets become available.

Finally, our analyses used freely available optical datasets. However, the problems associated with using optical imagery in the tropics, including the limited availability of cloud-free images over many regions and the rapid regeneration of tropical forest vegetation, remain major obstacles to pan-tropical assessments of tropical selective logging rates. Methods that integrate optical and radar dataset into a single algorithm would likely further improve the detection of tropical selective logging activities.

### 6. Conclusion

Loss and degradation of forests in the tropics has important implications for global climate change, local populations and biodiversity (Lewis et al., 2015). Methods to reliably map forest disturbances from selective logging would be a key contribution to quantifying the terrestrial portion of the carbon budget and the role of land-use change in tropical forests emissions (Baccini et al., 2017). In addition, reliable forest monitoring systems are actively sought after by tropical nations and conservation groups tasked with mitigating global climate change through improved forest management practices (GOFC-GOLD, 2016). Our results should stimulate further assessments of regional rates of low-intensity selective logging in tropical forests.

Our analysis, based on training Random Forest models with detailed records of tree removals, has demonstrated that Landsat data can be effective at detecting selective logging at much lower intensities than has previously been reported. To be successful, the input satellite data needs to be acquired within a few months of the logging, as the subtle signal caused by logging (and the more extensive disturbance associated with logging) is rapidly lost. Although we had less complete knowledge of logging activities at the Jari site, the algorithm developed at Jamari appeared to transfer successfully to this site (despite being 1500 km away). Hence there is reason to expect that it could be applied at much wider scales.

## Acknowledgements

We would like to thank *AMATA Brazil* for providing access to logging records for Jamari and *Jari Florestal* for logistical support. MGH was funded by the Grantham Centre for Sustainable Futures. JMBC was funded as part of NERC’s support of the National Centre for Earth Observation. FMF was CNPQ and NERC-funded (NE/P004512/1; PELD-RAS 441659/2016-0, respectively) and was funded by CAPES (BEX5528/13-5) and CNPq-PELD site 23 (403811/2012-0) during the long-term monitoring in Jari.

